# A Cellular Basis for Heightened Gut Sensitivity in Females

**DOI:** 10.1101/2025.05.23.654927

**Authors:** Archana Venkataraman, Eric E. Figueroa, Joel Castro, Fernanda M. Castro Navarro, Deepanshu Soota, Stuart M. Brierley, David Julius, Holly A. Ingraham

## Abstract

Visceral pain disorders, such as irritable bowel syndrome, exhibit a marked female prevalence. Enhanced signaling between enterochromaffin (EC) cells in the gut epithelium and mucosal sensory nerve fibers likely contributes to this sex bias. Here, we identify a novel estrogen-responsive paracrine pathway in which two enteroendocrine cell types, PYY-expressing L-cells and serotonergic EC cells, communicate to increase gut sensitivity in females. We demonstrate that ERα estrogen signaling upregulates the bacterial metabolite SCFA receptor *Olfr78* on colonic L-cells, increasing PYY release and their sensitivity to acetate. Elevated PYY acts on neighboring EC cells via NPY1R, thereby enhancing serotonin release and gut pain. We propose that hormonal fluctuations, in conjunction with internal (stress) or environmental (diet) factors, amplify this local estrogen-responsive colonic circuit, resulting in maladaptive gut sensitivity.

## MAIN TEXT

Chronic gastrointestinal pain conditions, such as irritable bowel syndrome (IBS), are more prevalent in women (*1, 2*) and can be exacerbated by fluctuating estrogen levels during the menstrual cycle and pregnancy (*3, 4*). While relevant targets of estrogen have not been identified, one possible site of action is the gut mucosa, where sensory nerve fibers communicate with epithelial cells to detect noxious stimuli and transmit nociceptive information to the spinal cord. Most epithelial cells that line the gut are enterocytes that primarily function to absorb nutrients, water, and electrolytes from the digestive tract. Interspersed among these enterocytes are rare, excitable enteroendocrine cells (EECs) that detect luminal contents (including nutrients, microbial metabolites, and ingested irritants) and release peptides and neurotransmitters to elicit an array of physiological responses. Serotonergic enterochromaffin (EC) cells are a distinct subtype of EECs that activate neighboring mucosal spinal afferents to elicit visceral pain (*5–7*). This EC cell-sensory nerve circuit shows heightened sensitivity in female mice (*5*), raising the possibility that it is a locus for hormone-dependent enhancement of visceral pain. Whether and how estrogen acts to modulate this peripheral pain circuit remains unclear.

The L-cell, another well-characterized EEC subtype, is primarily recognized for its role in sensing postprandial nutrients and secreting hormones such as glucagon-like peptide-1 (GLP-1) and peptide YY (PYY) to regulate insulin secretion, digestion, gastrointestinal motility, and absorption (*8*). While the sites of GLP-1 action are numerous, it is postulated to act locally through activation of GLP-1 receptors on EC cells to trigger the release of serotonin (*9*). However, unlike GLP-1, a role for PYY (specifically the cleaved PYY_3-36_ peptide) in promoting negative energy balance and appetite suppression (*10, 11*) is less clear (*12, 13*). When given exogenously, PYY_3-36_ results in significant GI discomfort in humans (*14, 15*) and food aversion in rodents (*16*). Whereas PYY_3-36_ selectively binds to NPY2 receptors in vagal afferents (*17*) and the brain (*18*), the larger PYY_1-36_ peptide additionally activates the NPY1 receptor (*19*), which in the gut are expressed by EC cells and spinal afferents (*9*). With EC cells emerging as the main detectors of noxious stimuli by the gut epithelium, coupling between PYY_1-36_ and NPY1R would presumably lead to visceral discomfort and hypersensitivity by increasing serotonin release, as originally hypothesized by Kojima and colleagues (*20*). However, given the intense focus on L-cells (and PYY) as nutrient sensors in the small intestine, few if any studies have explored paracrine coupling between L and EC cells in the colon and consequent effects on visceral pain. Here, we show that cellular crosstalk between L and EC cells leads to visceral hypersensitivity, revealing a role for PYY_1-36_ as a major nociceptive transmitter in the colon. This inter-enteroendocrine conduit is co-opted by estrogen via estrogen receptor alpha (ERα) on L-cells, revealing a mechanism to account for visceral hypersensitivity in females.

### ERα Signaling is Required for Visceral Sensitivity

We have previously shown that mucosal sensory afferents exhibit higher baseline activity in female mice compared to male mice (*5*). To determine if estrogen plays a role in this differential sensitivity, we compared afferent nerve fiber activity in ex vivo mucosal preparations (evMAR) from males versus intact or ovariectomized (OVX) females. Indeed, mechanical stimulation of the mucosa elicited higher responses in females compared to males, and this difference was greatly attenuated by OVX. (Fig. 1, A and B, and fig. S1, A to C). Similar results were obtained with evMAR preparations from mice (Na_V_1.8-ChR2) expressing channelrhodopsin in sensory afferents, where spiking was observed at substantially lower light intensities for females compared to males or OVX females (Fig. 1C and fig. S1D).

**Figure 1.**
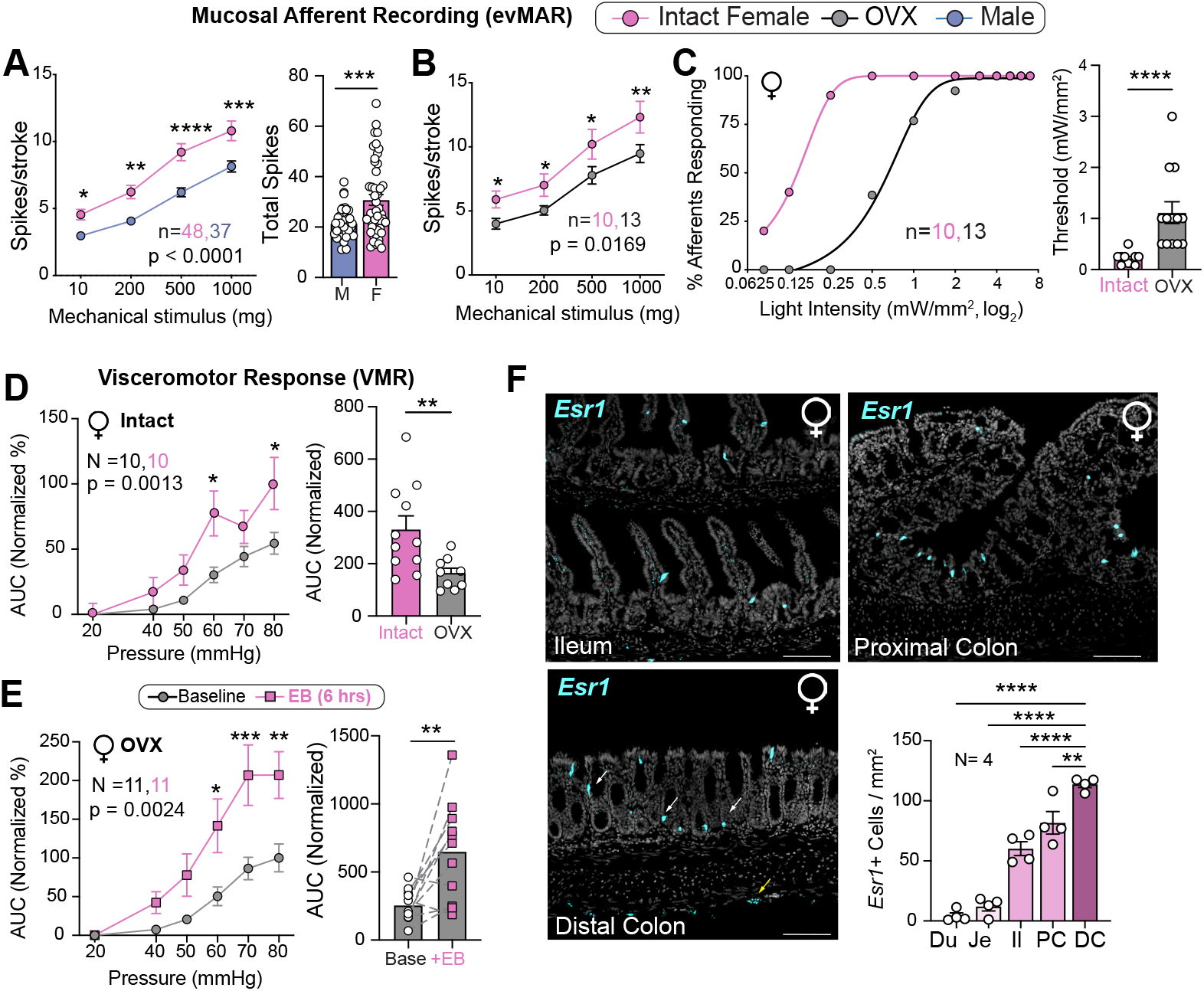
Estrogen Heightens Visceral Sensitivity in Females and is Enriched in the Distal Colon. **(A)** Mucosal afferent recordings (evMAR) from intact females and males to mucosal stroking (left). *n* = 37 to 48 afferents. Two-way repeated measures ANOVA followed by Bonferroni multiple comparisons test for each stimulus-response pair, \**P* < 0.05, \*\*\**P* < 0.001, \*\*\*\**P* < 0.0001. Total number of evMAR spikes (right). Unpaired t-test, \*\*\**P* < 0.001. (**B**) evMAR responses in intact and OVX females. *n* = 10 to 13 afferents. Two-way repeated measures ANOVA followed by Bonferroni multiple comparisons test, \**P* < 0.05, \*\**P* < 0.01. (**C**) Percentage of mucosal afferents responding at indicated light intensities (left), and activation thresholds of afferents (right) in intact and OVX *Na*_*V*_*1*.*8-ChR2* female mice. *n* = 10 to 13 afferents. Non-linear regression and the Mann-Whitney test were used, respectively, with \*\*\*\**P* < 0.0001 (**D**) VMR responses (left) and total area under the curve (AUC) (right) for intact and OVX females. *N* = 10 mice. Two-way repeated measures ANOVA followed by Šidák’s multiple comparisons test (left) and paired t-test (right). \**P* < 0.05, \*\**P* < 0.01. (**E**) VMR responses and total AUC in OVX female mice at baseline and 6 hours following estradiol benzoate (EB) injections (1µg/mouse). *N* = 11 mice. Two-way repeated measures ANOVA followed by Šidák’s multiple comparisons test (left), paired t-test (right). \**P* < 0.05, \*\**P* < 0.01, \*\*\**P* < 0.001. (**F**) *Esr1* transcripts visualized in the ileum, proximal, and distal colon, epithelial expression (white arrows) and neuronal expression (yellow arrows). Scale bars = 100 µm. Plot of *Esr1*+ cell density quantified from 15 fields per intestinal segment (Du=Duodenum, Je=Jejunum, Il=Ileum, PC=Proximal Colon, DC=Distal Colon). *N* = 4 mice. One-way ANOVA followed by Dunnett’s multiple comparisons test, \*\**P* < 0.01, \*\*\*\**P* < 0.0001. Data are presented as mean ± SEM.

Estrogen also had a marked effect on behavioral measures of visceral sensitivity as determined by assessing visceral motor responses (VMRs) to colorectal distension (CRD) (*21, 22*), akin to how pain is evaluated in patients with IBS (*23*). Responses in females were significantly blunted when estrogen was depleted by OVX (Fig. 1D). Remarkably, a single treatment of OVX females with estradiol benzoate (EB) reversed this drop, with maximum effects observed at 6 hours following EB injection (Fig. 1E and fig. S1E). Together, these results establish that estrogen, whether endogenous or exogenous, maintains a heightened state of gut sensitivity in females. In this regard, we were struck by our observation that the dominant transducer of estrogen, namely estrogen receptor alpha (ERα, encoded by the *Esr1* gene) is expressed at relatively low levels in proximal gut regions, but at much higher levels in a sparse population of cells within the colonic epithelium, particularly in the distal colon where visceral pain is most acutely sensed (Fig. 1F). To assess how loss of ERα affects gut function and visceral sensitivity, a *Vil1-Cre* driver was crossed to the floxed *Esr1*^*fl/fl*^ allele to eliminate ERα in the entire intestinal epithelium (Fig. 2A and fig. S2A). Gross metabolic parameters remained unchanged in mutant mice, including body weight and daily food intake (Fig. 2B and fig. S2, B and C). However, GI transit times and colonic motility were significantly accelerated, especially in females (Fig. 2, C and D), as predicted from known effects of estrogen on gut motility (*24*). Further, depleting intestinal ERα markedly lowered mucosal afferent sensitivity in females compared to males (Fig. 2E and fig. S2D). This lowered sensitivity was mirrored in VMR assays as *Esr1*^*Vil1-Cre*^ females were unresponsive to even the highest distension pressures (60-80 mmHg) compared to *Esr1*^*fl/fl*^ controls (Fig. 2F); circulating serotonin levels were also lower in mutant females (Fig. 2G). Especially striking was the inability of estrogen treatment to increase visceral sensitivity in *Esr1*^*Vil1-Cre*^ females, compared to their littermate controls (Fig. 2H, refer back to Fig 1E). These data establish that ERα is essential for estrogen-dependent enhancement of visceral sensitivity.

**Fig. 2.**
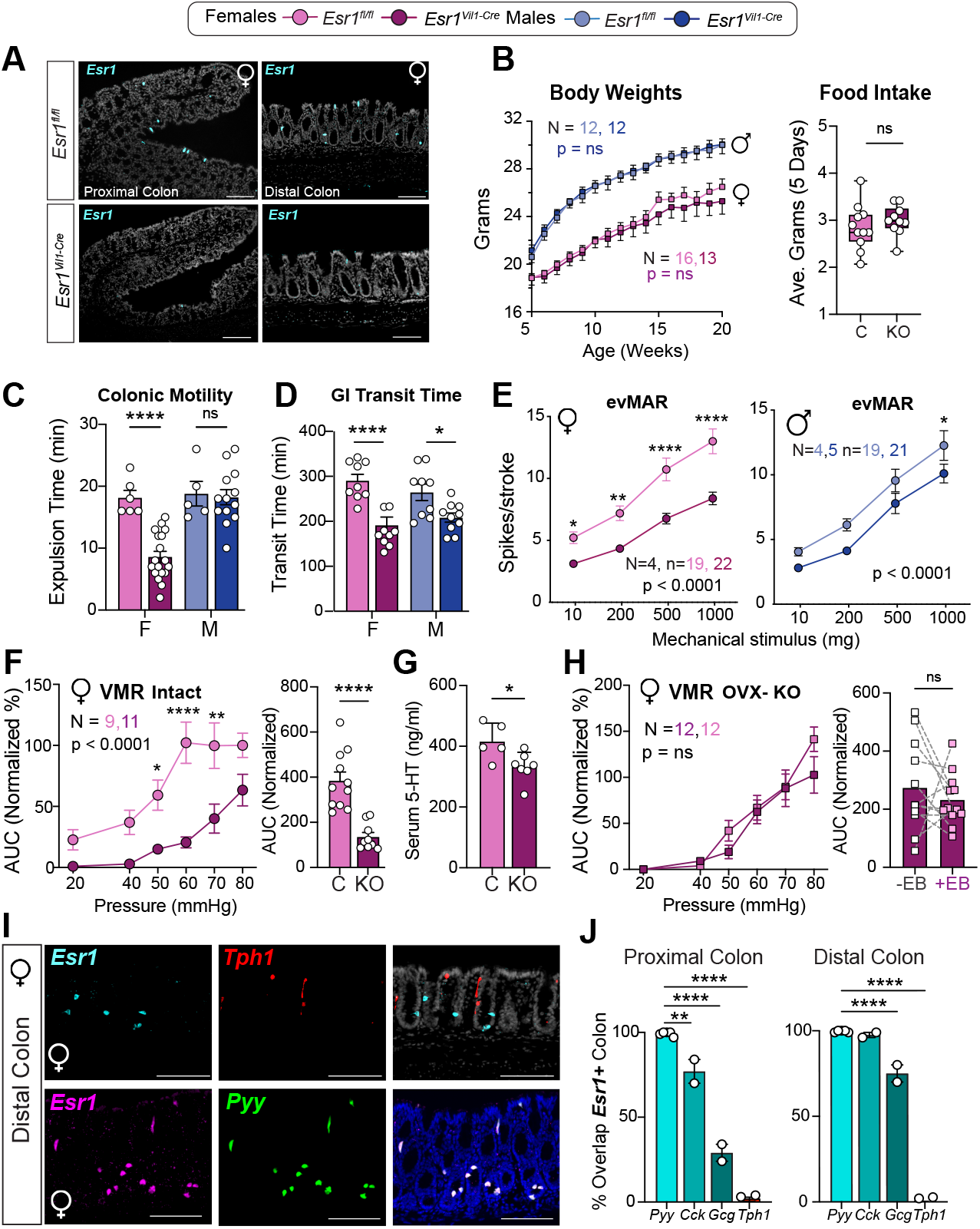
ERα Signaling Is Restricted to PYY^+^ L-Cells and Required for Gut Motility and Visceral Sensitivity. **(A)** *Esr1* expression in the proximal and distal colons of *Esr1*^*fl/fl*^ and *Esr1*^*Vil1-Cre*^ female mice. Scale bars = 100 µm. (**B**) Post-weaning body weights from 5 to 20 weeks in *Esr1*^*fl/fl*^ and *Esr1*^*Vil1-Cre*^ females and males (left); average food intake over a 5-day period in *Esr1*^*fl/fl*^ and *Esr1*^*Vil1-Cre*^ females (right). *N* = 11 to 16 mice. Two-way repeated measures ANOVA followed by Šidák’s multiple-comparisons test (left) and unpaired 2-tailed t-test (right), not significant (ns). (**C**) Colonic bead expulsion time in *Esr1*^*fl/fl*^ and *Esr1*^*Vil1-Cre*^ females (F) and males (M). *N* = 11 to 16 mice. Unpaired 2-tailed t-tests. \*\*\*\**P* < 0.0001, not significant (ns). (**D**) Gastrointestinal (GI) transit times in *Esr1*^*fl/fl*^ and *Esr1*^*Vil1-Cre*^ females (F) and males (M). *N* = 9 to 10 mice. Unpaired 2-tailed t-tests. \**P* < 0.05, \*\*\*\**P* < 0.0001. (**E**) evMAR responses in *Esr1*^*Vil1-Cre*^ and *Esr1*^*fl/fl*^ females and males. *n* = 19 to 22 afferents from 4 to 5 mice. Two-way repeated measures ANOVA followed by Šidák’s multiple-comparisons test. \**P* < 0.05, \*\**P* < 0.01, \*\*\*\**P* < 0.0001. (**F**) VMR responses and total AUC in intact *Esr1*^*fl/fl*^ controls and *Esr1*^*Vil1-Cre*^ females. *N* = 9 to 11 mice. Two-way repeated measures ANOVA followed by Šidák’s multiple-comparisons test (left) and unpaired 2-tailed t-test (right). \**P* < 0.05, \*\**P* < 0.01, \*\*\*\**P* < 0.0001. (**G**) Circulating 5-HT levels in *Esr1*^*fl/fl*^ and *Esr1*^*Vil1-Cre*^ females, unpaired 2-tailed t-test, \**P* < 0.05, *N* = 5 to 7 mice. (**H**) VMR data in OVX *Esr1*^*Vil1-Cre*^ females at baseline and following EB injections (1 µg/mouse). *N* = 9 to 10 mice. Two-way repeated measures ANOVA followed by Šidák’s multiple-comparisons test (left) and Wilcoxon matched-pairs signed-rank 2-tailed test (right), not significant (ns). **(I)** Images of female distal colons showing overlapping expression between *Esr1* and *Pyy* (bottom panel), but not *Tph1* transcripts (top panel). Scale bars = 100 µm. (**J**) Quantification of overlap between colonic *Esr1+* cells and *Pyy (N = 4), Cck, Gcg, and Tph1 (N = 2*) expressing enteroendocrine cells was examined across 15 fields per colonic segment. One-way ANOVA followed by Dunnett’s multiple comparisons test. \*\**P* < 0.01, \*\*\*\**P* < 0.0001. Data are presented as mean ± SEM.

Although we presumed ERα would reside in EC cells, given their role in driving visceral sensitivity, ERα was absent in EC cells (fig. S3A). While unexpected, this finding is consistent with the fact that very few estrogen-responsive genes were detected in EC cells (fig. S3B) and the number of EC cells was unchanged across the estrous cycle (fig. S3C). Instead, we found that ERα overlaps with 100% of *Pyy*- (and *Cck*) expressing cells, especially in the distal colon; overlap with *Gcg* that encodes GLP-1 was limited (Fig. 2, I and J, and fig. S3D).

### PYY-Induced Visceral Pain Elevated with Estrogen

Given the restricted expression of ERα in L-cells, we asked whether estrogen promotes visceral pain by increasing L-cell secretion of PYY and/or GLP-1, both of which have been proposed to enhance serotonin release via activation of their cognate G protein-coupled receptors on EC cells. In intact female mice, we found a marked increase in PYY levels peaking 6 hours post-estrogen treatment (Fig. 3A), whereas GLP-1 levels were largely unaffected (Fig. 3B) in this setting, even though estradiol has been shown to increase GLP-1 secretion (*25*). After first confirming expression of *Npy1r* on nociceptive EC cells (Fig. 3C), as reported by others (*9, 26*), we asked if PYY_1-36_ might act downstream of estrogen signaling in L-cells to enhance visceral sensitivity in female mice. Indeed, following the addition of PYY_1-36_ to the Na_V_1.8-ChR2 evMAR nerve-gut preparation, a prominent leftward shift was observed after light stimulation in OVX females (Fig. 3D and fig. S4, A and B). Moreover, PYY_1-36_ elicited increased afferent sensitivity to mechanical stimulation in OVX females (Fig. 3E and fig. S4, C and D). Of note, PYY failed to increase the higher baseline sensitivity in intact females (Fig. 3, F and G), but did increase visceral sensitivity in *Esr1*^*Vil1-Cre*^ KO mutant females and control males (fig. S5A, B). VMR responses to PYY_1-36_ were also significantly elevated in OVX control females or ERα KO mutant females (Fig. 3, H and I). These findings establish that PYY_1-36_ fully restores gut sensitivity following estrogen depletion in OVX females or after loss of estrogen signaling in the gut epithelium. PYY_1-36_ also increased afferent sensitivity in intact males, whereas GLP-1 failed to show any such effects in males or females (Fig. 3, J and K, and fig. S5C). Collectively, these data support the notion that the estrogen-induced PYY release from L-cells directly affects the EC-mucosal afferent circuit to increase visceral sensitivity.

**Fig. 3.**
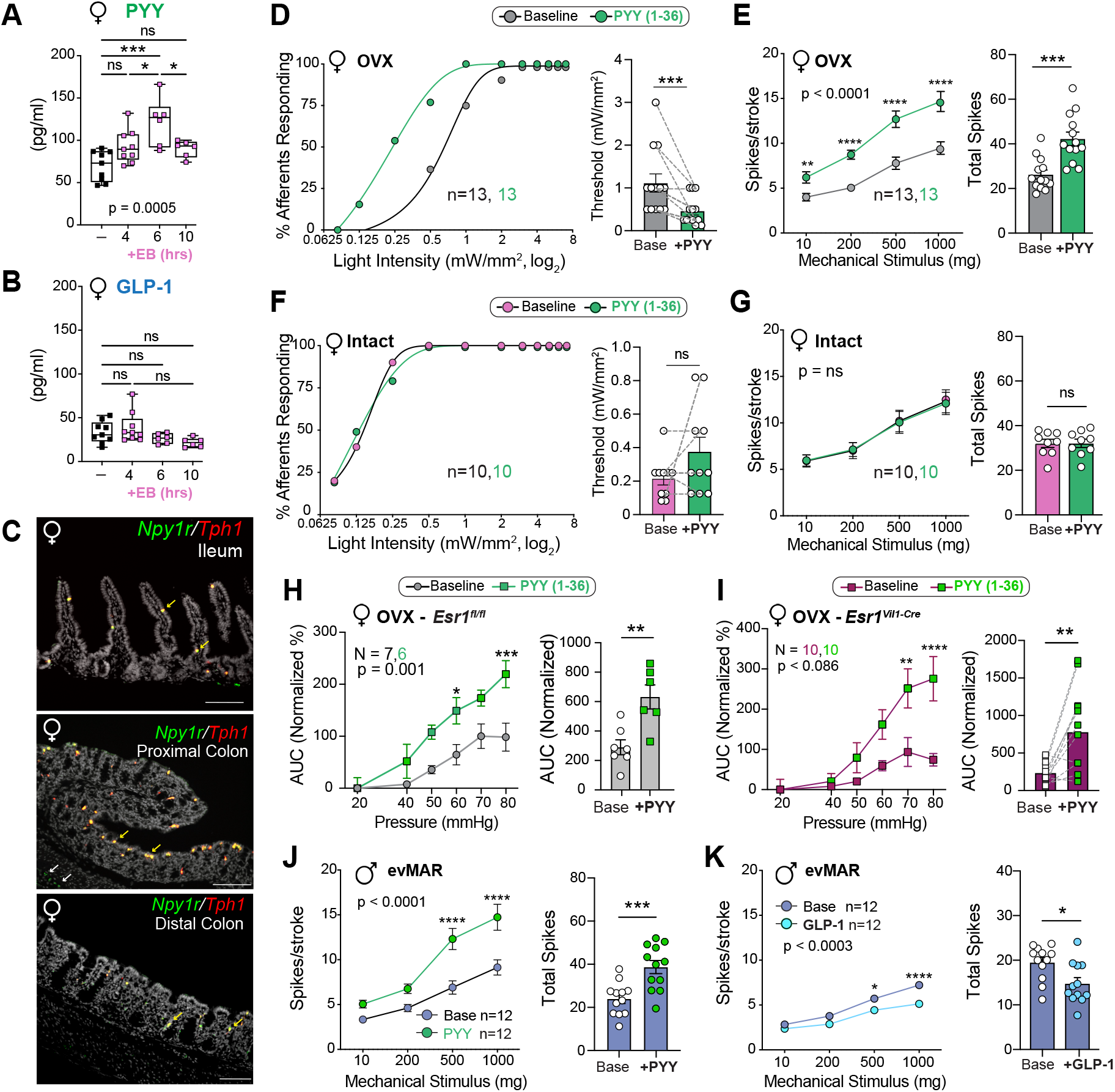
PYY_1-36_ but not GLP-1, Acts Downstream of ERα Signaling to Promote Visceral Sensitivity. **(A)** Circulating PYY levels and **(B)** circulating GLP-1 levels in female mice at baseline and following 4, 6, and 10 hours of EB (1 µg/ mouse) treatment. *N* = 6 to 10 mice. One-way ANOVA followed by Tukey’s multiple comparisons test. \**P* < 0.05, \*\*\**P* < 0.001, not significant (ns). (**C**) Representative images of *Npy1r* expression in *Tph1*+ EC cells visualized in the ileum, proximal, and distal colon, co-expression (white arrows) and neuronal expression (yellow arrows). Scale bars = 100 µm. (**D**) Percentage of mucosal afferents responding at indicated light intensities (left), and activation thresholds of afferents (right) in OVX *Na*_*V*_*1*.*8-ChR2* females at baseline and following PYY_1-36_ treatment (30 nM), *n* = 13 afferents. Non-linear regression (left) and Wilcoxon matched-pairs signed-rank 2-tailed test (right). \*\*\**P* < 0.001. (**E**) evMAR responses in OVX *Na*_*V*_*1*.*8-ChR2* females at baseline and following PYY_1-36_ treatment. *n* = 13 afferents. Two-way repeated measures ANOVA followed by Bonferroni multiple comparisons test (left) and unpaired 2-tailed t-test (right). \*\**P* < 0.01, \*\*\**P* < 0.001, \*\*\*\**P* < 0.0001. (**F**) Same as (D) but in intact *Na*_*V*_*1*.*8-ChR2* females at baseline following PYY_1-36_ treatment. *n* = 10 afferents. Non-linear regression (left) and Wilcoxon matched-pairs signed-rank 2-tailed test (right), not significant (ns). (**G**) Same as (E) but in intact *Na*_*V*_*1*.*8-ChR2* females at baseline and following PYY_1-36_ treatment. *n* = 10 afferents. Two-way repeated measures ANOVA followed by Bonferroni multiple comparisons test (left) and unpaired 2-tailed t-test (right), not significant (ns). (**H** and **I**) VMR responses and total AUC at baseline and after PYY ^1-36^ treatment (10 µg/kg) in OVX *Esr1fl/fl* control (H) and *Esr1*^*Vil1-Cre*^ KO (I) females. *N* = 6 to 10 mice. Two-way repeated measures ANOVA followed by Šidák’s multiple-comparisons test (left) and unpaired 2-tailed t-test in (H) (right) and paired 2-tailed t-test in (I) (right) \**P* < 0.05, \*\**P* < 0.01, \*\*\**P* < 0.001, \*\*\*\**P* < 0.0001. (**J** and **K**) evMAR responses in males at baseline and following PYY_1-36_ (J) or GLP-1_7-36_ (100 nM) (K) treatment. *n* = 12 afferents. Two-way repeated measures ANOVA followed by Bonferroni multiple comparisons test (left) and unpaired 2-tailed t-test (right). \**P* < 0.05, \*\*\**P* < 0.001, \*\*\*\**P* < 0.0001. Data are presented as mean ± SEM.

### A Local Paracrine Circuit Mediates Visceral Pain

To determine if PYY directly activates EC cells, we used live imaging of intestinal organoids engineered to express an EC cell-specific calcium reporter (*Polr2a-GCaMP5g*^*Tac1Cre*^). Robust PYY_1-36_ evoked signals were detected in male organoids (Fig. 4A), which were blocked by the NPY1R-specific antagonist, BIBO 3304 (Fig. 4B and fig. S6A). Similar responses were also detected in female organoids (Fig. 4, C and D, and fig. S6B). By contrast, PYY_3-36_ failed to elicit equivalent responses, again consistent with activation of the NPY1R subtype (Fig. 4E). Using a sniffer cell assay in which HEK293T cells express the gGRAB_5-HT3.0_ serotonin sensor (*27*), we found that PYY_1-36_ promotes robust serotonin release from EC cells. When sniffer cells (*6, 28*) were co-cultured with *Polr2a-GCaMP5g*^*Tac1Cre*^ organoids, PYY_1-36_ evoked serotonin release that was time locked with calcium responses in EC cells (Fig. 4F and G, and video S1). Together, these cellular assays demonstrate that L cell-derived PYY_1-36_ activates EC cells to elicit serotonin release, confirming previous studies measuring bulk transmitter levels (*20*). Blocking the NPY1R receptors in our *ex vivo* and *in vivo* assays yielded similar results. Indeed, following BIBO 3304 treatment, mucosal afferent responses in gut-nerve recordings from intact females were significantly reduced (Fig. 4H and fig. S6C). Moreover, estrogen-induced visceral sensitivity in OVX female mice was attenuated to baseline levels following BIBO 3304 treatment (Fig. 4I).

**Fig. 4.**
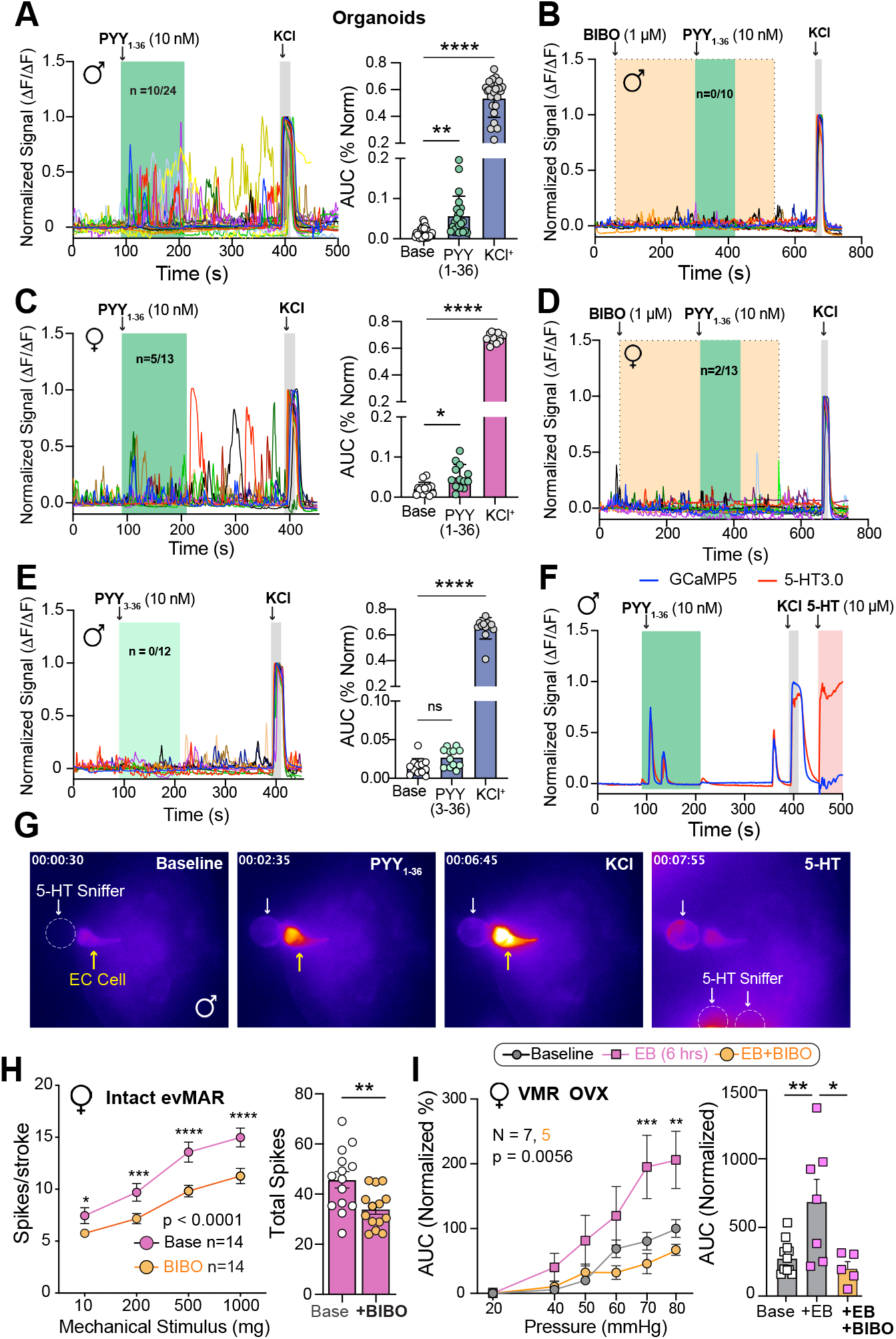
PYY_1-36_ Activates EC Cells to Promote Serotonin Release and Enhance Visceral Sensitivity via NPY1R. (A) PYY_1-36_ (10 nM) evoked calcium responses in EC cells within male intestinal organoids from *Tac1Cre;Polr2a(GCaMP5g-IRES-tdTomato*) mice. Responses are shown for each individual EC cell normalized to the maximal activation with KCl, with the number of responders indicated in the figure panel, *n* = 24 cells. Kruskal-Wallis test followed by Dunn’s multiple comparisons test, \*\**P* < 0.01, \*\*\*\**P* < 0.0001. (**B**) Effects on calcium responses in male organoids following BIBO 3304 (1 µM), an NPY1R antagonist, administered prior to PYY_1-36_, *n* = 10 cells. **(C)** Effects of PYY_1-36_ on female organoids as described in panel A. *n =* 13 cells. One-way ANOVA followed by Dunn’s multiple comparisons test, \**P* < 0.05, \*\*\*\**P* < 0.0001. (**D**) Effects of BIBO 3304 on female EC cells, *n* = 13 cells. (**E**) PYY_3-36_ effects on calcium responses in male organoids. *n* = 12 cells. One-way ANOVA followed by Dunnett’s multiple comparisons test for AUC data. \*\*\*\**P* < 0.0001, not significant (ns). (**F**) Representative recordings of gGRAB_5-HT3.0_-expressing HEK293T cells (red) and GCaMP5g-expressing EC cells in male organoids (blue) measuring 5-HT release and Ca2+ signals from EC cells in the sniffer experiment. (**G**) Representative frames from live imaging of gGRAB_5-HT3.0_ sniffer experiment with 5-HT sniffer cell (white arrow) next to a GCaMP5g+ EC cell (yellow arrow) in an intestinal organoid. A white-dashed circle outlines the sniffer cell in the far-left panel. (**H**) evMAR responses (left) and total spikes (right) in intact females at baseline and following BIBO 3304 treatment (1 µM). *n* = 14 afferents. Two-way repeated measures ANOVA followed by Bonferroni multiple comparisons test (left) and two-tailed, unpaired t-test (right). \**P* < 0.05, \*\**P* < 0.01, \*\*\**P* < 0.001, \*\*\*\**P* < 0.0001. (**I**) VMR responses and total AUC in OVX controls at baseline or treated with EB (1 µg) or EB + BIBO 3304 (1 mg/kg). *N* = 5 to 7 mice. Two-way repeated measures ANOVA followed by Šidák’s multiple comparisons test (left), and oneway ANOVA followed by Tukey’s multiple comparisons test (right). \**P* < 0.05, \*\**P* < 0.01, \*\*\**P* < 0.001. Data are presented as mean ± SEM.

To establish that PYY_1-36_ enhances mucosal afferent activity via serotonergic signaling, we administered PYY_1-36_ in combination with alosetron, a 5HT_3_R antagonist that is used clinically in women to mitigate IBS symptoms (*29*). Alosetron abrogated the PYY_1-36_-evoked sensitization of mechanical responses in gut-nerve recordings from OVX females (Fig. 5A and fig. S7, A and B). These findings were recapitulated *in vivo*, where alosetron, given 20 minutes before testing, effectively blocked all estrogen-induced visceral sensitivity in OVX females (Fig. 5B). Taken together, these data demonstrate that L cell-derived PYY_1-36_ acts locally to promote serotonin release and elicit gut pain by enhancing activity of the EC-mucosal afferent circuit.

**Fig. 5.**
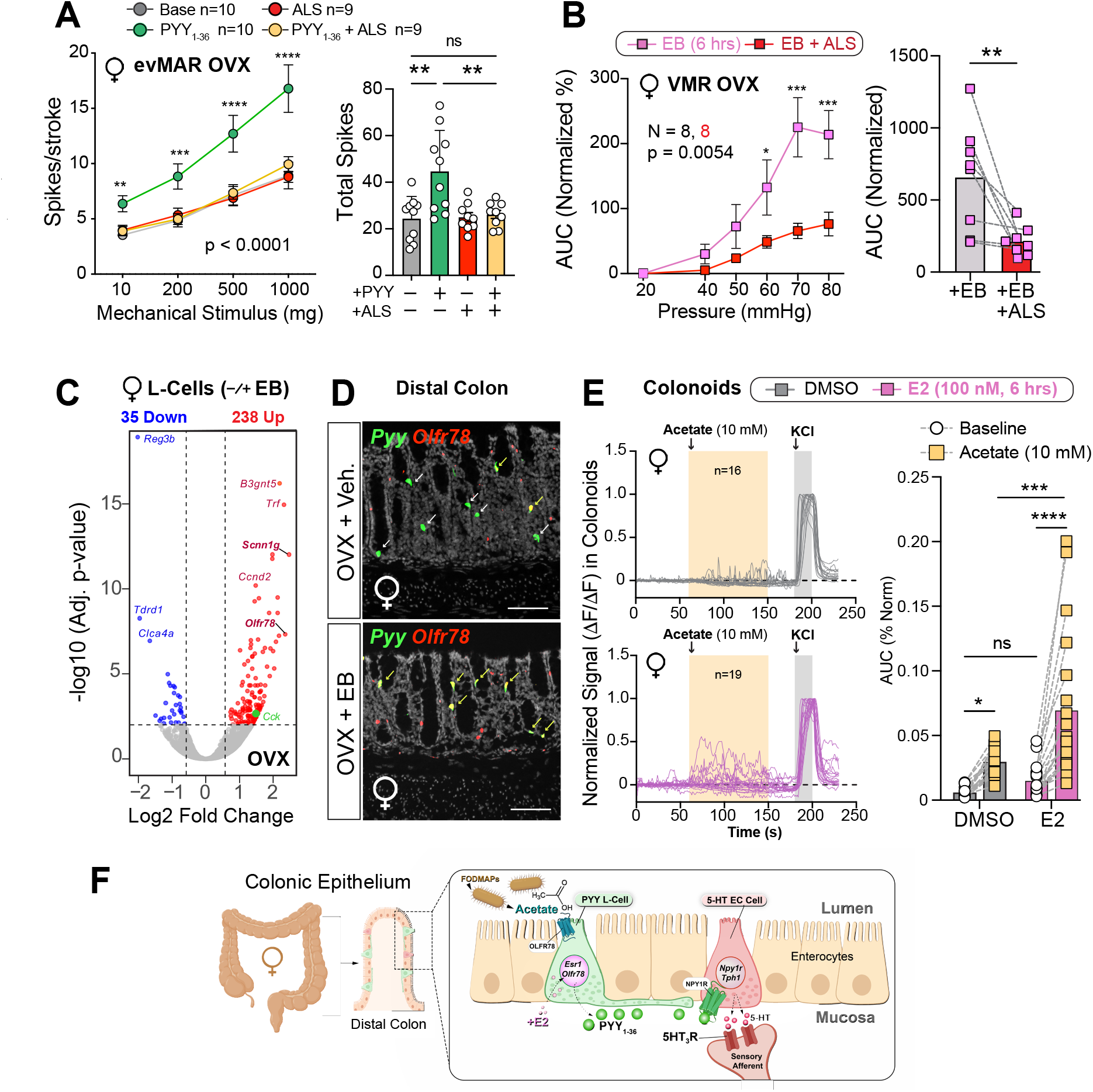
Estrogen Increases SCFA Detection in L-cells and Engages a Potent Gut Pain Pathway. (A) evMAR responses (left) and total spikes (right) in OVX females in four conditions-baseline, PYY_1-36_ (30 nM) treatment only, alosetron (ALS, 10 µM) treatment only, and PYY_1-36_ + ALS. *n* = 9-10 afferents. Two-way repeated measures ANOVA followed by Bonferroni multiple comparisons test (left) and one-way ANOVA followed by Tukey’s multiple comparisons test (right). \*\**P* < 0.01, \*\*\**P* < 0.001, \*\*\*\**P* < 0.0001, not significant (ns). (**B**) VMR responses and total AUC in OVX controls at baseline or treated with EB (1 µg/mouse) or EB + ALS (0.1 mg/kg). *N* = 5 to 7 mice. Two-way repeated measures ANOVA followed by Šidák’s multiple comparisons test (left), and one-tailed, paired t-test (right). \**P* < 0.05, \*\**P* < 0.01, \*\*\**P* < 0.001, \*\*\*\**P* < 0.0001. (**C**) Volcano plot showing changes in mRNA expression profile of L-cells in response to 4 hours of EB treatment, upregulated genes (red), downregulated genes (blue). (**D**) Representative images of distal colons showing *Olfr78* receptor expression in *Pyy* expressing L-cells from OVX mice treated with vehicle (OVX+Veh.) or estradiol benzoate (1 µg/mouse, OVX + EB). Scale bars = 100 µm. (**E**) Normalized fluorescent responses from L-cells expressing GCaMP5g at baseline (10 mM NaCl), and after application of acetate (10 mM), and high KCl in colonoids derived from *PyyCre;Polr2a(G-CaMP5g-IRES-tdTomato*) female mice preincubated with vehicle or E2 (100 nM) for 6 hours. n = 16 to 19 cells. Bar graphs for matched pairs of cells before and after treatment. Two-way ANOVA, mixed effects analyses. \**P* < 0.05, \*\*\**P* < 0.001, \*\*\*\**P* < 0.0001, not significant (ns). (**F**) Estrogen initiates paracrine signaling between L-cells and EC cells in the female colonic epithelium to affect visceral sensitivity. Estradiol acts through *Esr1* in L-cells to upregulate *Olfr78* expression, increasing its response to bacterial acetate and leading to PYY_1-36_ release. PYY_1-36_ then acts on NPY1R in enterochromaffin (EC) cells, resulting in activation of 5HT_3_R-expressing sensory nerve fibers to transduce pain information to the CNS. Data are presented as mean ± SEM.

### Estrogen Sensitizes L-Cell Response to SCFAs via *Olfr78*

Next, estrogen-responsive targets that might modulate visceral sensitivity were identified by profiling sorted labeled L-cells (*CckCre-tdTomato*) obtained from EB- or vehicle-treated OVX females. Multiple estrogen-sensitive targets, including known ERα targets (*Ccnd2, Cdc20*, and *Muc1*), were identified, with many more upregulated (248) than downregulated (35) (Fig. 5C and fig. S8A, tables S1-3). By contrast, a similar assessment of sorted EC cells revealed few if any estrogen-responsive genes (refer back to fig. S3B and table S4). The short-chain fatty acid (SCFA) receptor encoded by *Olfr78* emerged as a top hit, and its appearance in colonic L-cells after estrogen treatment was confirmed, as was the ENaC epithelial sodium channel subunit, *Scnn1g* (Fig. 5, C and D and fig. S8, B and C). By contrast, *Olfr78* was unchanged in sorted EC cells following estrogen treatment (refer back to fig. S3B). To assess whether upregulation of *Olfr78* is functionally relevant, female colonoids from *Polr2a-GCaMP5g*^*PyyCre*^ were preincubated with estradiol (E2) for 6 hours and then treated with acetate, the highest-affinity ligand for OLFR78 (*30–32*). Preincubating colonoids with estrogen significantly elevated L-cell responses to acetate (Fig. 5E). Collectively, our data suggest that the hormonal induction of *Olfr78* contributes to sensitization of the female gut through enhanced detection of bacterial metabolites.

## DISCUSSION

In this study, we have shown how estrogen signaling modulates paracrine interaction between two enteroendocrine cell types, elucidating a mechanism to explain heightened visceral sensitivity in females (Fig. 5F). PYY_1-36_ released from colonic L-cells initiates this cellular cascade by enhancing serotonergic tone in EC cells, leading to activation of near-by 5HT_3_R-expressing mucosal spinal afferents. Estrogen plays a crucial role in triggering these cellular events by 1) increasing PYY release from L-cells and 2) enhancing sensitivity of L-cells to bacterial metabolites through the upregulation of the SCFA receptor, *Olfr78*. Our findings also position EC cells as vital coincidence detectors in the gut-brain axis, corroborating earlier studies that paracrine interactions between L and EC cells increase serotonergic signaling, thereby affecting food intake (*9*), gut motility (*33*), and now visceral pain, as previously hypothesized (*20*).

Based on our pharmacological observations – namely, that estrogen enhances PYY release and that blocking NPY1R eliminates high visceral sensitivity in intact females or after restoring estrogen in OVX females– we place estrogen directly upstream of PYY-NPY1R signaling in EC cells. A local paracrine L-EC-sensory afferent signaling pathway, rather than direct action of PYY on NPY1Rs on spinal afferents (*34*), is further supported by the fact that the sensitizing effects of estrogen or PYY on visceral tone are abolished by antagonism of 5HT_3_R. Based on our inability to further sensitize visceral responses by PYY (Fig. 3, F and G) or SCFA (*5*) in intact females, we posit that when estrogen is fully engaged in intact females, PYY tone is already elevated, consistent with the higher levels of self-reported IBS in premenopausal women (*1, 35*). Our results raise the possibility that combining alosetron with a suitable NPY1R antagonist, possibly in conjunction with dietary reductions of short-chain fermentable carbohydrates or FODMAPs, known to elevate gut SCFAs and promote PYY release in an OLFR78-dependent manner (*32, 36*), might offer new therapeutic strategies to mitigate IBS or other forms of chronic visceral pain.

Our findings also help clarify conflicting views on the role of PYY in appetite suppression when viewed in the context of the colon and gut pain. We suggest that within the colon, PYY appears to be less important as a postprandial signal, and instead, functions as a major nociceptive hormone for eliciting pain and discomfort. The striking enrichment of ERα/PYY/*Olfr78*-positive cells (Fig. 1F and fig. 3SD) in the colon versus the small intestine (*32, 37, 38*) supports the notion that this estrogen-responsive molecular circuit is functionally positioned in the distal colon to sense and respond to environmental or endogenous noxious stimuli, especially in females. The profound changes observed in ERα−*Vil1Cre* mutant females in colonic motility and visceral hypersensitivity, but not energy balance, further support the differential action of PYY in modulating nociception versus metabolic responses at these locales.

We predict that fluctuating estrogen levels would impact the crosstalk between L and EC cells, and two periods in the female life cycle come to mind – surges in estrogen during the follicular and luteal phases, and the meteoric rise in estrogen during late stages of pregnancy. Clearly, increasing sensitivity to environmental stimuli in the mother’s gut would be protective for developing progeny. Taken together with recent studies that have begun to define molecular cues that promote villus expansion in the maternal gut (*39, 40*), our study illuminates yet another essential adaptation to enhance progeny wellbeing, namely, heightened surveillance of ingested nutrients. Identifying this protective signaling mechanism brings us closer to understanding how hormones and diet, when coupled with stress and inflammatory events, could become maladaptive, leading to chronic visceral pain.

## SUPPLEMENTAL MATERIALS AND METHODS

### Ethics

All experiments in the study were approved and performed following the Institutional Animal Care and Use Committees (IACUC) guidelines set by University of California, San Francisco, and South Australian Health and Medical Research Institute (SAHMRI), the National Institutes of Health Guide for Care and Use of Laboratory Animals, and recommendations of the International Association for the Study of Pain. Adult mice (>8 weeks old) were housed on a 12-hour light/ dark cycle in a humidity and temperature-controlled housing facility with ad libitum access to food and sterile water unless otherwise noted for experiments.

### Mice

*Esr1*^*fl/fl*^ mice were maintained in the laboratory on a mixed CD-1;129P2 background, as previously described (*41*). Hemizygous *Vil1-Cre* mice (Jackson Laboratories, Strain #021504) were bred with homozygous *Esr1*^*fl/fl*^ mice to achieve intestine epithelial cell-specific knockout of *Esr1* in *Esr1*^*Vil1-Cre*^ mice. For optogenetic evMAR experiments, *Scn10a*-Cre mice (gift from W. Imlach, Monash University, Australia) were crossed with *ChR2*-floxed mice (Jackson Laboratories, Strain #012569) to get Na_V_1.8-ChR2 mice. *Scn10a* (Na_V_1.8) is expressed by >95% of colonic afferents (*42*), and conditional expression of light-sensitive channelrhodopsin-2 (ChR2) in these neurons allowed targeted optogenetic activation of colonic mucosal afferents but not EECs. *Cck-Cre* mice (Jackson Laboratories, Strain #012706) or *Pyy-Cre* mice (*43*) (gift from D. Bohórquez, Duke University, USA) were crossed with reporter *R26LSL-tdTomato* mice (Jackson Laboratories, Strain #007914) to label colonic *Esr1*+ cells. GCaMP imaging in organoids used *Tac1-Cre* mice (Jackson Laboratory, Strain #021877) or *Pyy-Cre* crossed with *Polr2a(GCaMP5g-IRES-tdTomato*) reporter mice (gift from L. Jan, UCSF; Jackson Laboratory, Strain #024477). Cre- and Flp-dependent reporter mouse line, *RC::FLTG* (Jackson Laboratory, Strain #026932), was crossed with the transgenic *Pet1-Flp* and *Tac1-Cre* lines as previously described (*5*). This allowed us to label EC cells exclusively with the tdTomato reporter in the resulting EC^FLTG^ animals *RC::-FLTG; Pet1-Flp::Tac1-Cre. Cre*-negative littermate controls were used in experiments where possible. Cohorts of ovariectomized (OVX) mice were created by surgical removal of ovaries in 8-12-week-old females; all in vivo, ex vivo assays or collection of colonic tissue were performed 2-3 weeks after ovariectomy.

### Colon-pelvic nerve preparation for flat-sheet single fiber mucosal afferent recordings

Male and female C57BL/6J, *Esr1*^*fl/fl*^ or *Esr1*^*Vil1-Cre*^ mice were humanely killed by CO_2_ inhalation. The colon and rectum with attached pelvic nerves were removed, and recordings from mucosa afferents were performed as previously described (*5, 44*). Briefly, the colon was removed and pinned flat, mucosal side up, in a specialized organ bath. The colonic compartment was superfused with a modified Krebs solution (in mM: 117.9 NaCl, 4.7 KCl, 25 NaHCO^3^, 1.3 NaH^2^PO^4^, 1.2 MgSO^4^ (H^2^O)^7^, 2.5 CaCl^2^, 11.1 D-glucose), bubbled with carbogen (95% O_2_, 5% CO_2_) at a temperature of 34ºC. All preparations contained the L-type calcium channel antagonist nifedipine (1 µM) to suppress smooth muscle activity and the prostaglandin synthesis inhibitor indomethacin (3 µM) to suppress potential inhibitory actions of endogenous prostaglandins. The pelvic nerve bundle was extended into a paraffin-filled recording compartment in which finely dissected strands were laid onto a mirror, and a single fiber was placed on the platinum recording electrode. Action potentials generated by mechanical stimuli to the colon’s receptive field pass through the fibers into a differential amplifier, filtered, sampled (20 kHz) using a 1401 interface (CED, Cambridge, UK), and stored on a PC for off-line analysis. Categorization of the properties of afferents followed our previously published classification system (*44*). Receptive fields were identified by systematically stroking the mucosal surface with a still brush to activate all subtypes of mechanoreceptors. In short, pelvic mucosal afferents respond to delicate mucosal stroking (10 mg von Frey hairs; vfh) but not circular stretch (5 g). Stimulus-response functions were then constructed by assessing the total number of action potentials generated in response to mucosal stroking with 10, 200, 500, and 1000 mg vfhs. For C57/BL6J mice, PYY (30 nM) was applied in combination with the DPP IV inhibitor K579 (1 µM) or GLP-1_7-36_ (100 nM) to the mucosal epithelium for 5 min via a small metal ring placed over the receptive field of interest, and mechanosensitivity was re-tested. In some studies, co-application of PYY and K579 was preceded by the application of the 5-HT3R antagonist alosetron (10 µM, incubated for 10 minutes). In separate studies, BIBO 3304 (1 µM) was applied to the mucosal epithelium for 10 min via a small metal ring placed over the receptive field of interest, and mechanosensitivity was re-tested. For *Esr1*^*fl/fl*^ and *Esr1*^*Vil1-Cre*^ mice, PYY (30 nM) in combination with the DPP IV inhibitor K579 (1 µM) was applied to the mucosal epithelium for 5 min via a small metal ring placed over the receptive field of interest, and mechanosensitivity was re-tested.

### Optogenetic stimulation of colonic mucosal afferents

As *Scn10a* (Na_V_1.8) is expressed by >95% of colonic afferents (*42*) but not by ECs (*6*), we crossed *Scn10a*-Cre mice with ChR2-floxed mice to produce *Na*_*V*_*1*.*8-ChR2* mice, allowing testing of colonic afferent responses to both mechanical and light stimuli, as previously described (*5, 28*). Colonic recordings were prepared from *Na*_*V*_*1*.*8-ChR2* mice using methods described above. Colonic mucosal afferents were identified by their responsiveness to fine mucosal stroking (10 mg vfh), but not to circular stretch (5 g). We then recorded the action potentials generated by stroking the colonic mucosa with calibrated vfh (10, 200, 500, and 1000 mg), applied for 2 seconds with a 10-second interval between stimuli. Afferents were left to rest for 10 minutes before optogenetic stimulation. We then recorded the action potentials generated by stimulating the receptive field of mucosal afferents with continuous exposures of increasing light (470 nm) intensities, ranging from 0.082-7 mW. Each light exposure was applied for 2 seconds with a 10-second interval between exposures. Light was delivered to a small section of the colon (~3 mm^2^) covering the afferent’s receptive field, via a High Power Fiber-Coupled LED Light Source (model BLS-FCS-0470-10) and Multimode Fiber Patchcords (Numerical aperture: 0.39 NA, Core size: 400 µm. Catalog #FPC-0400-39-025MA-BP, Mightex, Pleasanton, CA 94566, US). Following baseline mechanical and optogenetic stimulation, PYY (30 nM) in combination with the DPP IV inhibitor K579 (1 µM) was applied to the mucosal epithelium via a small metal ring placed over the receptive field of interest. Five minutes later, afferent sensitivity to mechanical and optogenetic stimulation was re-tested, in the presence of PYY and K579. Action potentials were analyzed offline using the Spike 2 wavemark function and discriminated as single units based on a distinguishable waveform, amplitude, and duration (CED, Cambridge, UK). Optogenetic stimulation was operated by BLS-SERIES software (BLS-SA04-US).

### *In vivo* assessment of colonic pain

Visceromotor responses (VMR) to colorectal distensions (CRD) were assessed using electromyography (EMG) in fully awake animals as described previously (*5*). To allow for repeated measurements across time on the same animal, a wireless transmitter (ETA-F10; Data Sciences International, New Brighton, MN, USA) was placed subcutaneously with the leads inserted into the abdominal musculature. Animals were housed individually post-surgery and allowed to recover for at least 10 days before VMR tests. Under brief anesthesia, the distal colon was cleared of its contents, and a lubricated 2 cm long balloon was gently inserted into the colorectum up to 0.25 cm beyond the anal verge. Once the balloon catheter was secured to the base of the tail, the animal was moved to a restrainer and allowed to recover from anesthesia for 15 minutes. The balloon catheter was then connected to a barostat (Isobar 3, G&J Electronics, Willowdale, Canada) for graded and pressure-controlled delivery of the balloon distension sequence. The distensions were 20s-long with 2-min intervals and applied in the following sequence: 20, 40, 50, 60, 70, and 80 mmHg.

For OVX females receiving estradiol benzoate (EB) treatment, animals were injected subcutaneously with sesame oil (VEH; Sigma, Catalog #S3547) or 1 µg of EB (Cayman Chemical, Catalog #10006487) per mouse. VMR experiments were performed 6 hours after EB injections, coincident with peak PYY release. Repeated VMR measurements in the same animal were spaced at least 5 days apart. For animals treated with PYY_1-36_ (10 µg/kg s.c., Bachem, Catalog #4031137) in combination with DPP4 inhibitor (K579, 1mg/ kg i.c., Sigma-Aldrich Catalog #317642), or with alosetron (0.1 mg/kg s.c., Sigma-Aldrich Catalog #SML0346), or with BIBO 3304 (1mg/kg i.c., Tocris Bioscience, Catalog #2412), drugs were administered 20 minutes before the start of the distension sequence. EMG recordings in response to the distensions were relayed to Ponemah Software (Data Sciences International) data acquisition system and analyzed offline using SpikEB Software (CED). The magnitude of VMR responses to each distension pressure was quantified by computing the area under the curve (AUC) during the distension (20 seconds) corrected for baseline EMG activity 20 seconds immediately before the distension. All VMR values were then normalized to the average VMR response of control animals in each experimental group to the maximum distension (i.e., average control VMR response to 80 mmHg set at 100%). Total AUC was calculated as the summation of VMR responses across all distension pressures for each animal.

### Circulating PYY, GLP-1 and 5-HT Measurements

Mice were anesthetized with 1.5% isoflurane and submandibular venous blood was collected in heparin-coated tubes with 10% protease inhibitor cocktail (v/v) containing 5000 kIU/ml Aprotinin (Research Products International, Catalog #A20575), 1.2 mg/ml EDTA, and 0.1 nmol/l Diprotin A (Sigma-Aldrich, Catalog #I9759). Plasma was isolated by centrifugation at 1,500 g for 10 minutes at 4°C and stored at −80°C until further analysis. For total PYY and GLP-1 measurements, a customized MSD U-PLEX platform (MesoScale Discovery, Catalog # K152ACM) was used. For 5-HT measurements, blood was collected in microcapillary tubes (SAFE-T-FILL) with clot activator. After centrifugation at 10,000 rcf for 10 minutes, serum was collected and stored at −20 °C until further analysis. Serotonin levels were quantified using an ELISA kit (Eagle Biosciences) and Spectra-Max iD5 microplate reader (Molecular Devices).

### Fluorescence In Situ Hybridization (FISH) and Immunofluorescence

Intestinal segments from 8–12-week-old female and male mice were cryosectioned to 10 µm thickness, and RNA-FISH was performed using the RNAscope Multiplex Fluorescent v2 Assay (Cat. No. 323110, Advanced Cell Diagnostics) according to the manufacturer’s instructions. RNAScope probes against Esr1 (Cat. No. 478201), Pyy (Cat. No. 420681), Cck (Cat. No. 402271), Tph1 (Cat. No. 318701), Npy1r (Cat. No. 427021), and Gcg (Cat. No. 400601) were used. TSA Plus Fluorescence dyes Fluorescein (Cat. No. NEL741001KT), Cy3 (Cat. No. NEL744001KT), or Cy5 (Cat. No. NEL745001KT) from Akoya Biosciences were diluted in TSA buffer. All sections were counterstained with DAPI nuclear stain (Cat. No. 320858).

Confocal images were acquired at the UCSF Nikon Imaging Center using a Nikon Ti2 microscope with a Crest LFOV spinning disk and MicroManager v.2.0 gamma. Images were processed and quantified using ImageJ FIJI software. To quantify *Esr1* expression across the intestine, 15 images per segment per animal were captured on the confocal microscope at 20X magnification. The number of *Esr1+* cells were manually quantified, normalized to the total area of intestinal epithelium, and then averaged across biological replicates.

For immunofluorescence, 5 µm and 10 µm intestinal cryosections were stained overnight with primary antibodies at 4°C at the indicated dilutions. Antibodies used were against ERα (EMD Millipore, polyclonal rabbit, 1:500 dilution), and serotonin (Immunostar, polyclonal goat, 1:3000). Sections were then labeled with species-appropriate secondary Alexa-Fluor-coupled antibodies (Invitrogen, 1:500 dilution) for detection.

### Food Intake Studies

*Esr1*^*fl/fl*^ and *Esr1*^*Vil1-Cre*^ mice were singly housed with access to a FED3 device set to free-feeding mode for ad libitum food access (*45*). All animals used in the study learned to successfully retrieve 20-mg pellets (BioServ, F0071) from the dispenser within 48 hours of acclimation. Daily body weights and food intake measurements were recorded for the next five days. Animals were fasted for 24 hours on day 8 and then re-feeding responses were assessed the following day.

### Gastrointestinal transit studies

Mice were fasted for 6 hours before the start of the transit experiments with *ad libitum* access to water. For measurements of gastrointestinal transit time (GITT), a 300 µl solution of 6% (w/v) carmine red in 0.5% methylcellulose (Sigma, C1022) was administered by orogastric gavage. Total GI transit time was calculated from the time of administration to the appearance of the first red stool pellet. For colonic transit measurements, mice were briefly anesthetized with isoflurane, and a 3 mm diameter glass bead was inserted in the colon to a depth of 2 cm. The time to bead expulsion was then recorded.

### Quantitative PCR analysis

Colonic segments were washed with ice-cold PBS and subjected to mechanical dissociation and EDTA incubation to isolate epithelial cells. Cells were then suspended in Trizol solution (Ambion). RNA was isolated through RNeasy columns (Qiagen) and converted to cDNA using the SuperScript III First-Strand Synthesis System (ThermoFisher). Quantitative RT-PCR was performed using SYBR Green Select Master Mix (Thermo Fisher). Primer sequences are listed in table S5.

### Preparing and culturing of intestinal organoids

Adult male and female *Tac1-Cre;Polr2a*^*GCaMP5g-IRES-tdTomato*^ mice were used to generate intestinal organoids as previously reported (*28*). The upper jejunum was specifically used to avoid ectopic *Tac1Cre* expression in the lower intestine. Organoids were maintained and passaged every 6 days in organoid growth medium (advanced Dulbecco’s modified Eagle’s medium–F12 supplemented with penicillin–streptomycin, 10 mM HEPES, Glutamax, B27 (Thermo Fisher Scientific), 1 mM N-acetylcysteine (Sigma), 50 ng ml^−1^ mouse recombinant epidermal growth factor (Thermo Fisher Scientific), R-spondin 1 (10% final volume) and 100 ng ml^−1^ murine Noggin (Peprotech)). R-spondin 1-conditioned media was obtained from R-spondin 1-expressing HEK293T cells (Sigma) maintained in DMEM, 20% FBS, 1% penicillin-streptomycin, and 125 μg ml^−1^ zeocin (Thermo Fisher Scientific) at 37 °C, 5% CO2.

Adult female *PYY-Cre;Polr2a*^*GCaMP5g-IRES-tdTomato*^ mice were used to generate colonoids from the distal colon as previously reported (*46*). Colonoids were maintained and passaged every 7 days in colonoid growth medium (advanced Dulbecco’s modified Eagle’s medium–F12 without phenol-red supplemented with 1% penicillin–streptomycin, 10 mM HEPES, 1X Glutamax, B27 Plus (Thermo Fisher Scientific), 1 mM N-acetylcysteine (Sigma), and Wnt-3A, R-spondin 3, and noggin-conditioned (L-WRN) media (50% final volume). L-WRN cells (ATCC; Cat.# CRL-3276) were maintained in DMEM without phenol red, 20% Charcoal/Dextran-treated FBS, 1% penicillin–streptomycin, 0.5 mg mL^−1^ G-418 and 0.5 mg ml^−1^ hygromycin B. G-418 and hygromycin B were removed to produce Wnt-3A, R-spondin 3, and noggin.

### GCaMP imaging of intestinal and colonic organoids

All pharmacological reagents for intestinal organoids as listed in table S5, were delivered by local perfusion at the following concentrations: PYY_1-36_ (10 nM), PYY_3-36_ (10 nM), the DPP4 inhibitor, sitagliptin (200 nM), 5-HT (10 µM), E2 (100 nM), sodium acetate (10 mM) and KCl (70 mM).

Five days after passage *Tac1-Cre;Polr2a*^*GCaMP5g-IRES-tdTomato*^ small intestine organoids, and seven days after passage *Pyy-Cre;Polr2a*^*GCaMP5g-IRES-tdTomato*^ colonoids were removed from Matrigel (Corning) and broken up by sequential trituration using a 1,000 μl pipette tip, followed by a 200 μl pipette tip. The organoid fragments were seeded onto Cell-Tak (Corning)-coated coverslips and placed in a recording chamber containing Ringer’s solution (140 mM NaCl, 5 mM KCl, 2 mM CaCl_2_, 2 mM MgCl_2_, 10 mM Glucose and 10 mM HEPES-Na, pH 7.4) or Ringers-Krebs-HEPES solution (119 mM NaCl, 5 mM KCl, 25 mM HEPES-Na, 1.2 mM MgSO_4_ 1.2 mM KH^2^PO^4^ and 2 mM CaCl^2^ (pH 7.35; 10 mM glucose added before use). EC cells or L-cells were identified by td-Tomato expression. GCaMP imaging was performed with an upright microscope equipped with a Grasshopper 3 (FLIR) camera and a Lambda 421 optical beam combiner (Sutter Instrument). Organoids were maintained under a constant laminar flow of Ringer’s or Ringer’s-Krebs-HEPES solution applied by a pressure-driven microperfusion system (SmartSquirt, Automate Scientific). Acquired images were analyzed using Fiji software (NIH). ROIs were drawn around individual EC cells, and ΔF/(F_max_-F_0_) values were calculated and normalized to a positive control (e.g., 70 mM KCl). Cells with responses greater than 2 S.D. from baseline AUC mean were considered responders to pharmacological treatment.

### Biosensor experiments

HEK293T cells stably expressing gGRAB_5HT3.0_-IRES-mCherryCAAX were maintained in DMEM, 20% FCS, 1% penicillin–streptomycin and 2 μg ml^−1^ puromycin (Thermo Fisher Scientific) at 37 °C, 5% CO_2_ Cells were dissociated with TrypLE™ Express (GIBCO™), washed once with Ringer’s solution and plated on top of intestinal organoids. Individual HEK293T cells were carefully lifted from coverslips and positioned at least 5 μm away from a tdTomato+ EC cell using a glass pipette. Imaging was performed using an inverted microscope equipped with a Grasshopper 3 camera (FLIR) run using the Micro-Manager software (v.2.0) and a Lambda 721 optical beam combiner (Sutter Instrument). The entire area of each biosensor cell was used to calculate ΔF/ (F_max_-F_0_) values. At the end of each recording, gGRAB_5-HT3.0_ was fully activated with 10 μM serotonin. Bath solution was maintained under a constant laminar flow of Ringer’s solution applied by a pressure-driven microperfusion system (SmartSquirt, Automate Scientific).

Pharmacological reagents (dissolved in Ringer’s solution as described above) were delivered by local perfusion for all organoid and colonoid imaging studies. Acquired images were analyzed using Fiji software (NIH). ROIs were drawn around individual EC cells, L-cells, and biosensor cells, and the resulting signals were calculated and normalized to a positive control ΔF/(F_max_-F_0_), including 70 mM KCl or 10 µM 5-HT.

### FACS isolation and bulk RNA sequencing

To isolate epithelial cells, colonic tissue was cut into ~3 cm segments and filleted. The tissue was subsequently incubated in 10 mL of cold PBS with 30 mM EDTA and 1.5 mM dithiothreitol (DTT) at 4°C for 45 minutes on a tumbling rotator. The tissue was transferred to 5 mL of pre-warmed PBS with 30 mM EDTA and incubated at 37°C for 8 minutes. During incubation, the tissue was shaken every minute. The dissociated epithelium was washed with 10 mL DPBS with 10% fetal bovine serum (FBS) and digested in 10 mL of HBSS with 0.5U/mL dispase II (Sigma) and 5 μM Y-27632, and 0.2 mg/mL DNaseI (Sigma) at 37°C for 14 minutes, with vigorous shaking at 2-minute intervals. The cells were then washed with 10 mL HBSS with 10% FBS and 5 μM Y-27632, and 0.2 mg/mL DNaseI. Digested and washed cells were resuspended in 500 μL of sorting media (DMEM/F12, 10 mM HEPES, 1x B27 supplement, 5 μM Y-27632, and 0.2 mg/ mL DNaseI) supplemented with antibodies as described. Stained cells were washed twice with DMEM/F12 with 10 mM HEPES, 0.5% FBS, and 5 μM Y-27632, and 0.2 mg/mL DNaseI and subsequently filtered through 70μm and 40μm strainers. Filtered cells were reconstituted in sorting media and kept on ice until sorting.

L-cells were isolated from the colons of 10–14-week-old OVX *Cck-Cre;R26LSL-tdTomato* female mice treated with sesame oil (VEH) or estradiol benzoate (EB) 4 hours prior. Colonic epithelial cells were isolated using the EDTA/ dispase method and mechanical disruption as described above. Single-cell suspensions were stained with EpCAM antibody (BioLegend) to select for epithelial cells and DAPI (Sigma-Aldrich) to exclude dead cells. Approximately 1% of total EpCAM+ epithelial cells were tdTomato+ and sorted using BD FACSAria FUSION Cell Sorter (UCSF Laboratory for Cell Analysis). Approximately 7,000 L-cells from both treatment groups (OVX + VEH and OVX + EB) were collected into TRIzol LS (Thermo Fisher). EC cells were isolated from the colons 10-14 OVX EC^FLTG^ females. At least 500 EC cells per sample were processed similarly from OVX EC^FLTG^ mice treated with VEH or EB. A complete list of antibodies used for FACS purification of EECs is provided in table S6.

RNA extraction, library preparation, and sequencing of the sorted cells were performed using Genewiz next-generation sequencing services (South Plainfield, NJ, USA). PolyA cDNA libraries were sequenced on the Illumina Nextera XT platform, generating 150 bp paired-end reads. Approximately 50M reads were obtained per sample of tdT+ L-cells, and ~15-20M reads per sample of tdT+ EC cells. Raw RNA-seq reads were processed using the Galaxy server (v4.9). The reads were aligned to the mm10 mouse reference genome using RNA STAR (*47*). The resulting BAM files were subsequently used as input for featureCounts (*48*) to quantify mapped reads in an unstranded manner. Raw counts from different datasets were then imported into the DESeq2 pipeline (v2.11.40.8) (*49*) to identify differentially expressed genes and generate normalized counts using default thresholds.

### Data availability

RNA-seq datasets have been deposited at GEO (https://www.ncbi.nlm.nih.gov/geo/) under the SuperSeries accession number GEO: (Provided with Updates).

### Statistics, blinding, and randomization

All statistical tests are described in each figure legend, with additional details provided for all panels in table S7. Prior to statistical analyses, all data were first analyzed for normal distribution using Kolmogorov-Smirnov or Shapiro-Wilk tests, with the appropriate statistical test and post hoc test specified in each figure legend. All behavior, GCaMP, and nerve recordings were performed by blinded experimenters, with blinding codes revealed post-analysis to allow for statistical comparisons. VMR data were normalized with the average maximal response taken as 100%. Optogenetic mucosal afferent data are expressed as the threshold for action potential activation (mW/mm^2^). Unless otherwise stated, data are presented as mean ± SEM and analyzed using Prism software (version 9.50, GraphPad, San Diego, CA, USA). Differences were considered statistically significant at *p* < 0.05. *N* = number of animals, and n = number of afferents/cells/organoids. Genetically modified mice and control littermates were randomly allocated from different cages of male and female mice for analyses. Group sizes of males and females were used to achieve sufficient power for statistical significance for all measurements, and after the genotypes of mice were unblinded for VMR, evMAR recordings, behavioral, and cellular physiological assays. For all experiments involving drug vs. vehicle, animal selection was randomized.

## ACKNOWLEDGMENTS

We thank the Garvan Institute, Australia, for genotyping services, the Preclinical, Imaging and Research Laboratories (PIRL, SAHMRI) for the use of their small animal facility, the UCSF Nikon Imaging Core for use of their confocal imaging facilities, and R. Sarah Elmes for assistance in FACS analyses. We thank Ms. Callie Goodman for for help with the initial characterization of ERα (*Esr1*^*Vil1Cre*^) mutant mice, Drs. James Bayrer, Zsofia Torok, Koki Touhara, and Sarah Mohr, for many helpful suggestions and critical comments during these studies.

This work was supported by NIH Training Grant T32 DK007418 and NIGMS K12GM081266-17 to E.E.F; NIDDK R01DK135714 to H.A.I., D.J., and S.M.B.; R35 NS105038 to D.J.; National Health and Medical Research Council of Australia (NHMRC) Investigator Leadership Grant APP2008727 to S.M.B., an NHMRC Development Grant APP2014250 to S.M.B, and an NHMRC Ideas Grant APP1181448 to J.C.

## DATA AVAILABILITY STATEMENT

All data generated or analyzed during this study will be included in the published article (and its supplementary information files).

## AUTHOR CONTRIBUTIONS

Conceptualization: AV, EEF, DJ, HAI, SMB

Methodology: AV, EEF, JC, FMCN, DS

Investigation: AV, EEF, JC, FMCN

Visualization: AV, EEF, JC, FMCN, DS, HAI, SMB

Funding acquisition: EEF, HAI, DJ, SMB, JC

Supervision: HAI, DJ, SMB

Writing – original draft: AV, EEF, DJ, HAI

Writing – review & editing: AV, EEF, DJ, HAI, JC, SMB

## SUPPLEMENTAL DATA FIGURES AND TABLES

Data will be provided upon request.

